# Single cell Iso-Sequencing enables rapid genome annotation for scRNAseq analysis

**DOI:** 10.1101/2021.09.27.461747

**Authors:** Hope M. Healey, Susan Bassham, William A. Cresko

## Abstract

Single cell RNA sequencing (scRNAseq) is a powerful technique that continues to expand across various biological applications. However, incomplete 3′ UTR annotations in less developed or non-model systems can impede single cell analysis resulting in genes that are partially or completely uncounted. Performing scRNAseq with incomplete 3′ UTR annotations can impede the identification of cell identities and gene expression patterns and lead to erroneous biological inferences. We demonstrate that performing single cell isoform sequencing (ScISOr-Seq) in tandem with scRNAseq can rapidly improve 3′ UTR annotations. Using threespine stickleback fish (*Gasterosteus aculeatus*), we show that gene models resulting from a minimal embryonic ScISOr-Seq dataset retained 26.1% greater scRNAseq reads than gene models from Ensembl alone. Furthermore, pooling our ScISOr-Seq isoforms with a previously published adult bulk Iso-Seq dataset from stickleback, and merging the annotation with the Ensembl gene models, resulted in a marginal improvement (+0.8%) over the ScISOr-Seq only dataset. In addition, isoforms identified by ScISOr-Seq included thousands of new splicing variants. The improved gene models obtained using ScISOr-Seq lead to successful identification of cell types and increased the reads identified of many genes in our scRNAseq stickleback dataset. Our work illuminates ScISOr-Seq as a cost-effective and efficient mechanism to rapidly annotate genomes for scRNAseq.

## INTRODUCTION

Single cell RNA sequencing (scRNAseq) is a revolutionary technique in biology that provides expression information from tissues and embryos (E. Shapiro, Biezuner, and Linnarsson 2013). By barcoding RNA from individual cells directly from dissociated samples, scRNAseq allows for post hoc analysis of cell types and can be used to ascertain novel cell populations, explore developmental trajectories, and define gene regulatory networks (Luecken and Theis 2019).

To maximize the utility of scRNAseq datasets, however, 3′ UTRs must be annotated for several reasons. ScRNAseq captures transcripts through poly(A) tails leading to a 3′ bias in coverage (Hwang, Lee, and Bang 2018). Partially annotated genes may be represented in a dataset with lower read counts, leading to erroneous conclusions regarding their magnitude of expression across cell types. In addition, downstream scRNAseq analysis clusters cell types through the determination of a covariance structure across highly variable genes (Stuart et al. 2019). The systematic absence of such genes can lead to inferential errors in multivariate analyses and obscure biological reality.

The annotations of even intensively studied models, such as mice and zebrafish, continue to be improved (Gupta et al. 2018; Lawson et al. 2020). Most other organismal genomes are even less well annotated. To facilitate a broader utility of scRNAseq requires more efficient and methods for 3′ UTR annotation. Recently, full length, single-molecule isoform sequencing (Iso-Seq) has been used to improve genome annotations (Ali, Thorgaard, and Salem 2021; Beiki et al. 2019; Kuo et al. 2017, 2020). PacBio’s Iso-seq has been further adapted to use the 10x Genomics platform for scRNA barcoding (ScISOr-Seq) to track cell type specific isoform expression (Gupta et al. 2018; Zheng et al. 2020).

Here we show that ScISOr-Seq in the context of a scRNAseq experiment allows rapid 3′UTR annotation in threespine stickleback fish (*Gasterosteus aculeatus*). This fish has long been a focus of study in behavior, ecology and evolution (Bell and Foster 1994; Colosimo et al. 2004; Cresko et al. 2004, 2007; Hohenlohe et al. 2010; Reid, Bell, and Veeramah 2021; M. D. Shapiro et al. 2004), and is now a nascent system for biomedical research (E A Beck et al. 2020; Emily A. Beck et al. 2021; Fuess et al. 2021; Gardell et al. 2017; Miller et al. 2007; Small et al. 2017). Although stickleback have a well assembled genome, it’s 3′ UTR annotations are incomplete which limits scRNAseq’s utility. We demonstrate that a single PacBio SMRT cell of ScISOr-Seq data is sufficient to significantly improve the stickleback annotations to an extent on par with zebrafish for the purpose of scRNAseq analysis at this stage. Our findings demonstrate that ScISOr-Seq will be a useful tool to efficiently improve genome annotations for scRNAseq in many organisms.

## RESULTS AND DISCUSSION

### ScISOr-Seq captured novel isoforms and improved 3′UTR

The ScISOr-Seq reads were classified with SQANTI3 (Tardaguila et al. 2018) using Ensembl gene models to describe isoforms based on how well they match splice variants (Figure 1A). The most common structural category (45.15% of isoforms) is “novel not in catalog” containing at least one new splicing site relative to the existing annotation (Figure 1A and 1B). While 17.42% of collapsed isoforms were “full splice matches” (FSM), only 16.7% of these isoforms matched the annotation. Therefore, roughly 3% of the unique isoforms matched the existing gene annotations, the rest improve existing or added gene models. Confirming that 3′ UTRs were poorly annotated, 46.2% FSM had alternative 3′ ends and 27.4% had alternative 3′ and 5’ ends (Figure 1C). Therefore, the existing stickleback annotation - in addition to having incomplete 3′ UTRs - is also missing many splice variants. Our work indicates that additional ScISOr-Seq or bulk Iso-Seq is necessary to capture these variants as well as prune erroneous transcript models from the Ensembl annotation.

**Figure 1:**
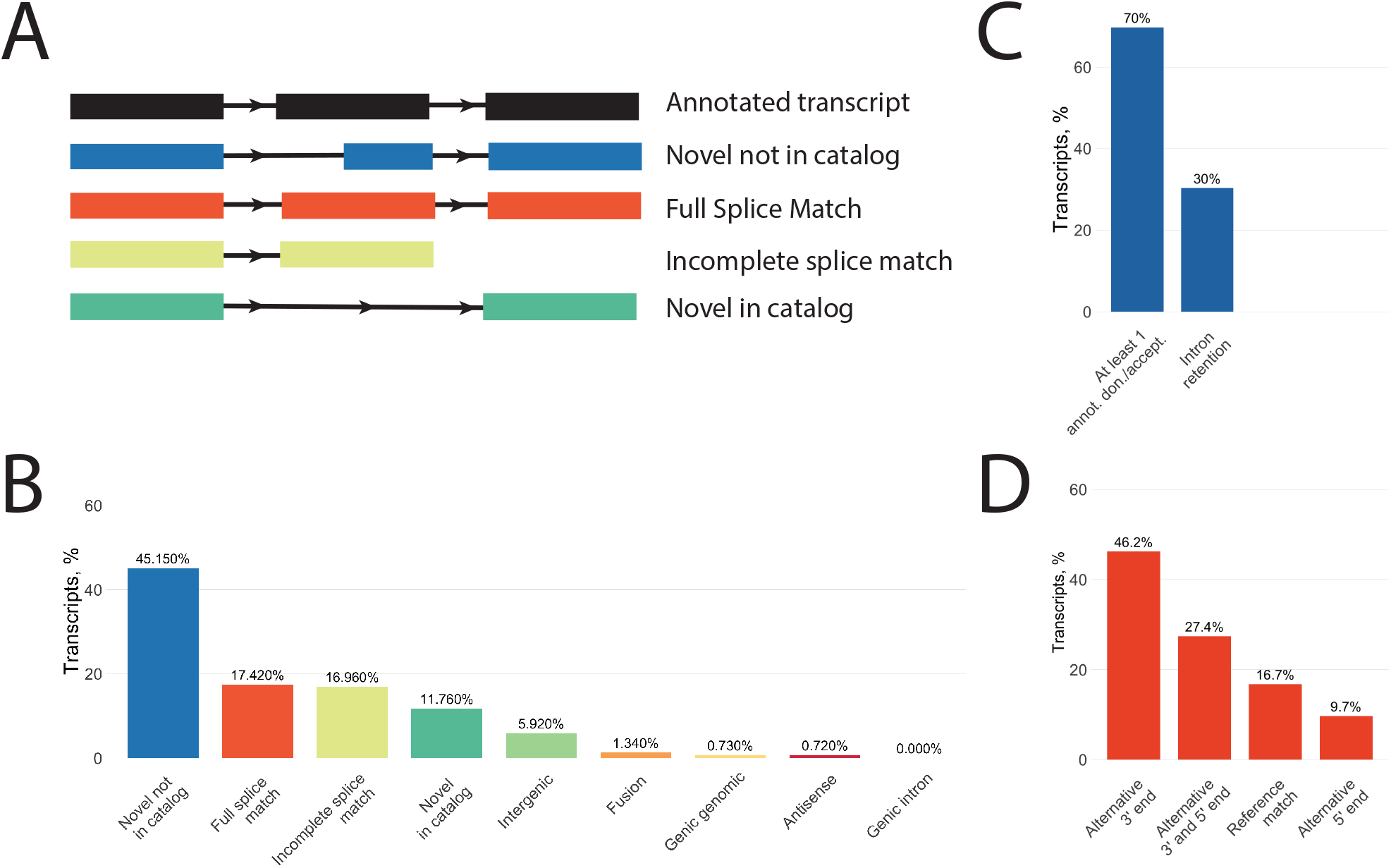
SQANTI3 classification of ScISOr-Seq data revealed that the majority of isoforms were previously unannotated in the stickleback genome. A) SQANTI3 categorizes isoforms based on their match to the reference, the major categories are shown here. B) the distribution of ScISOr-Seq isoforms in the major SQANTI3 classes illustrating that the bulk of isoforms are novel not in catalog. C) within the novel not in catalog category, isoforms are largely coming from cases where there is at least one new splicing donor or acceptor with some coming from cases of intron retention (for example, *nkx2.3* isoforms from Figure 2). D) the majority of isoforms that are full splice matches have at least an alternative 3′ site while few are complete reference matches.

### ScISOr-Seq improved the number of reads retained for scRNAseq

We initially tested how well the existing stickleback gene models from Ensembl (BROAD S1, 104.1 database version) would capture scRNAseq reads. Strikingly, less than 50% of reads were retained for downstream scRNAseq analysis (Figure 2B; Supplemental file 1). Using such an incomplete annotation for scRNAseq would likely result in erroneous interpretation of gene expression patterns for genes lacking 3′ UTR annotations.

**Figure 2:**
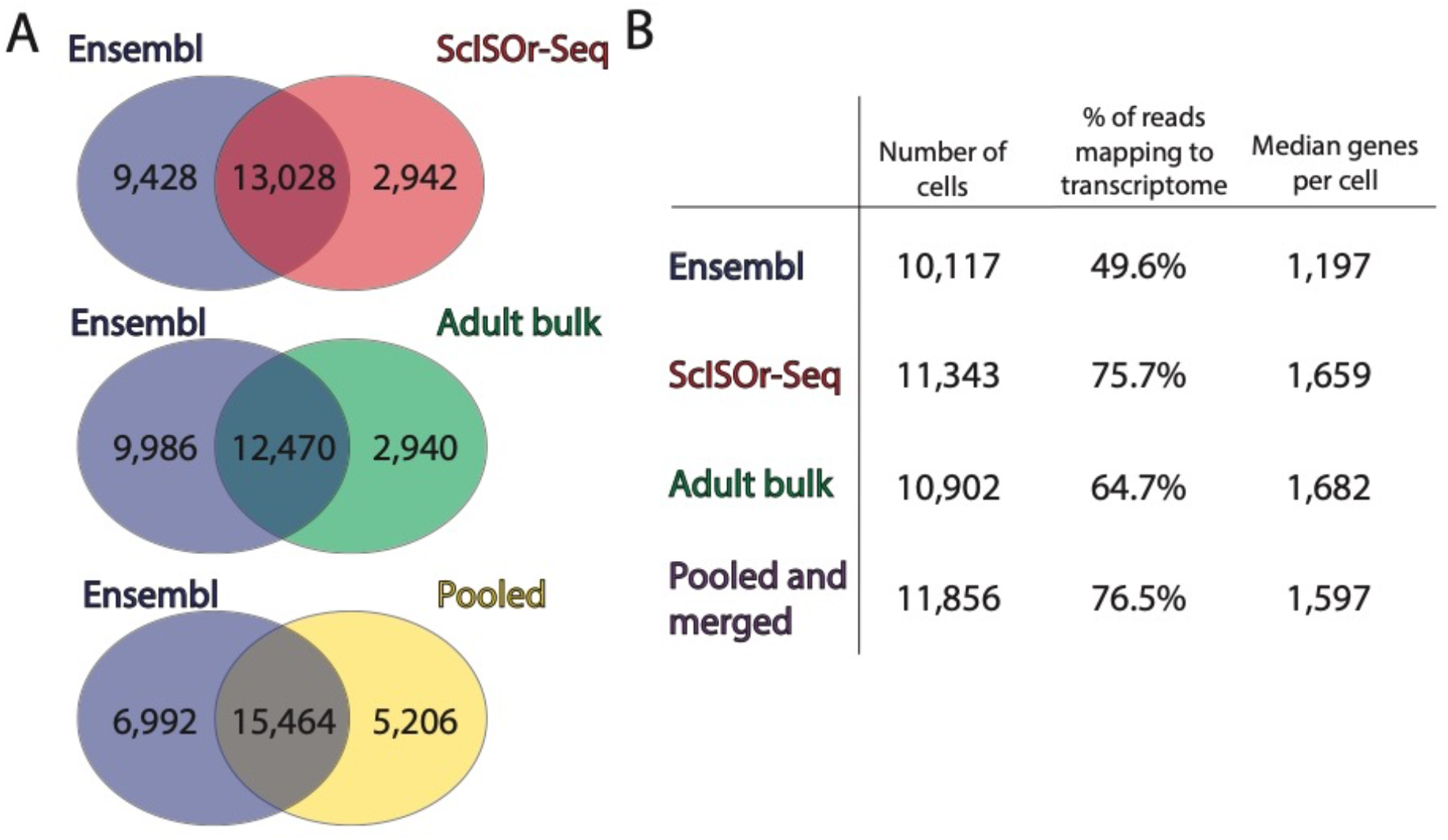
ScISOr-Seq and bulk Iso-Seq captured a limited number of annotated transcripts, but both resulted in an increase in reads mapping to the transcriptome. Ultimately, pooling this data and merging with the ensembl annotation resulted in the largest gains in reads mapped. A) the number of genes in common and unique to each Iso-Seq read category is compared to the ensembl genome. B) results from cell ranger runs using the different genome annotation files illustrating the overall improvements by using Iso-Seq data particularly the ScISOr-Seq gene models.

Next, we used ScISOr-sequencing data to generate new gene models and tested how well these new models would capture scRNAseq reads (Supplemental file 2). Using the ScISOr-Seq dataset, 13,028 previously annotated genes and 2,942 novel genes were identified (Figure 2A). The ScISOr-Seq annotations lead to a notable 26.1% increase in reads retained from scRNAseq compared to the Ensembl gene models alone (Figure 2B; Supplemental file 1). The alignment of scRNAseq reads with existing Ensembl and new ScISOr-Seq gene models illustrates that ScISOr-Seq models retain greater numbers of reads due to improved annotation of 3′ UTRs (Figure 3; Figure S1A and S1B). This result compares favorably with current research in the much better studied zebrafish model. Farnsworth, Saunders, and Miller (2020) completed their zebrafish atlas with 80% of reads retained in 5dpf (days post fertilization) fish. After Lawson et al. (2020) improved the zebrafish transcriptome, they noted a 4% increase in reads retained that lead to an increase of 2,257 cells and 8 clusters using the same 5 dpf atlas (Farnsworth, Saunders, and Miller 2020). The similarity in number of retained reads in our dataset with those published in zebrafish indicates that these improvements in stickleback are sufficient for scRNAseq.

**Figure 3:**
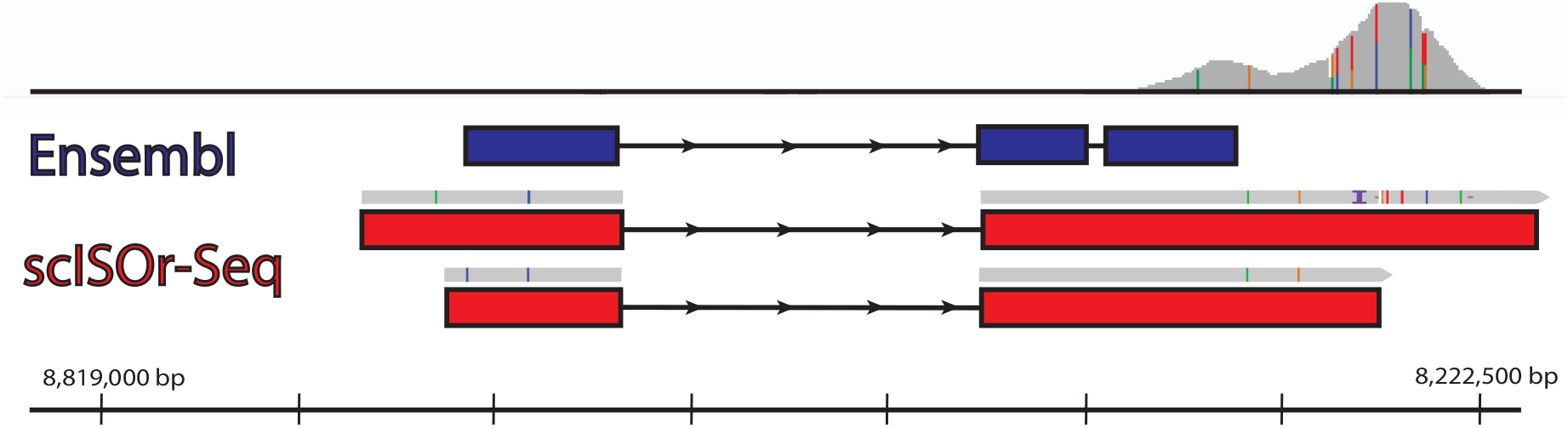
ScISOr-Seq improved gene models by extending the 3′ UTRs illustrated here by the two models for nkx2.3. scRNAseq reads are shown at the top in grey. The existing ensembl model is shown in dark blue. Notably, this model does not overlap with the majority of the scRNAseq reads. The ScISOr-Seq reads (in grey) are shown above the respective gene models (in red) that they generated. Colored lines on the ScISOr-Seq and scRNAseq reads indicate a different base pair in the read than the stickleback reference genome (BROAD S1, 104.1 database version). These ScISOr-Seq gene models capture all of the scRNAseq reads.

Recently, Naftaly, Pau, and White (2021) published a large bulk analysis of adult stickleback Iso-Seq data from 16 PacBio SMRT cells, which we reanalyzed to create an annotation to test if comparably more scRNAseq reads are retained (Supplemental file 3). Although we identified similar overall annotated and novel genes, the bulk Iso-Seq annotation from adult stickleback captured 11% fewer reads than our embryonic ScISOr-Seq annotation (Figure 2A and 2B; Supplemental file 1). These results are likely due to differential expression of transcripts between the adult and embryonic samples, highlight that even for species with previous Iso-Seq libraries additional sequencing may be needed for scRNAseq analysis in other developmental stages or tissue types.

Since both our ScISOr-Seq and the bulk Iso-Seq gtf files improved scRNAseq reads retained, Because of the improvements with ScISOr-Seq and bulk Iso-Seq we pooled them to create a comprehensive annotation and also merged this pooled dataset with the Ensembl annotations. We retained all transcripts and united them under a single gene model for scRNAseq analysis (Supplemental file 4). This new annotation contained 6,992 genes unique to Ensembl, 15,464 genes in both datasets, and 5,206 genes unique to the pooled dataset (Figure 2A). Although pooling and merging added 16 SMRT cells of Iso-Seq data as well as the full suite of Ensembl annotations, this effort only improved the proportion of reads retained by +0.8% beyond the annotation created by just our single ScISOr-Seq sample (Figure 2B; Supplemental file 1). Therefore, the gene models originating from ScISOr-Seq data alone could be used for scRNAseq analysis if a system lacks a prior annotation.

### ScISOr-Seq enhances biological interpretation of cell clusters

Using the pooled (ScISOr-Seq and bulk Iso-Seq) annotation that was merged with Ensembl annotation, we proceeded with scRNAseq analysis (Supplemental file 5) to test whether biologically meaningful cell clusters would be created. We clustered cells by their transcriptional profiles into cell identities and analyzed the results with Seurat (Stuart et al. 2019). We identified 30 clusters of cells with 38 principal components (Figure 4A). We putatively annotated cell types via distinguishing sets of marker genes based on zebrafish literature (Farnsworth, Saunders, and Miller 2020; D. G. Howe et al. 2013; Wagner et al. 2018).

**Figure 4:**
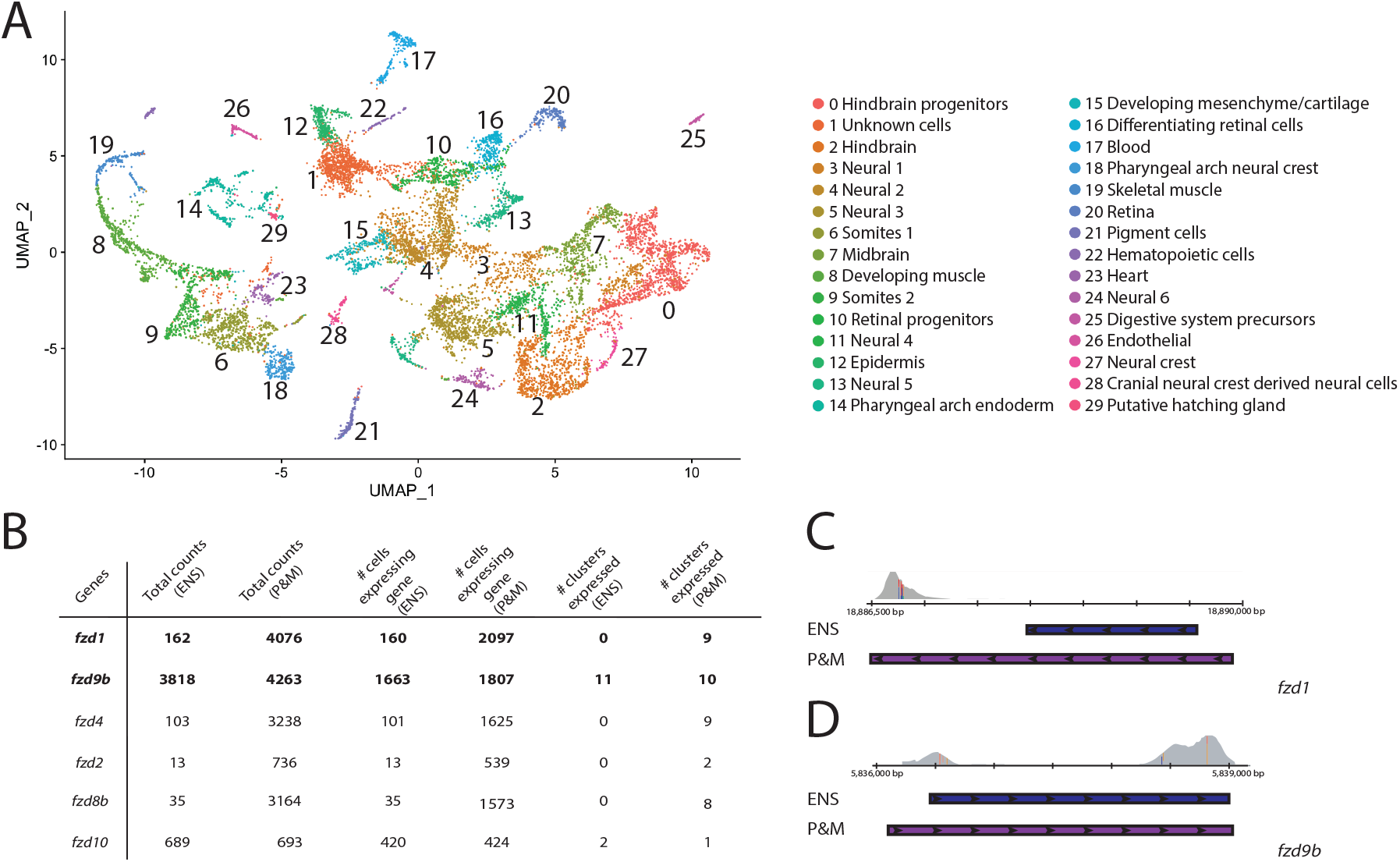
Using the updated gene models from our merged and pooled dataset, we identified expected cell types in 70hpf stickleback scRNAseq data and illustrated how annotations improvements are relevant for analysis. A) the clustering and cell identities of the scRNAseq data is illustrated on the UMAP. B) frizzled gene family expression compared between the ensembl annotation scRNAseq analysis and the pooled and merged analysis using total raw counts, number of cells expressing each gene, and the number of clusters where over 10% of cells are expressing each gene. C) *fzd1* gene model is greatly extended in the pooled and merged annotation (purple) compared to the ensembl model (blue) which allows for much greater counting of reads (grey). D) fzd9b had minor changes in the pooled and merged model (purple), a longer 5’ UTR, in comparison with the ensembl annotation (blue) which led to minimal increases in reads (grey) counted. The scRNAseq read pileups in C and D have colored lines on base pairs that are different from the stickleback reference genome (BROAD S1, 104.1 database version).

Although zebrafish and stickleback diverged ~ 229.9 m.y.a. (K. L. Howe et al. 2021), we identified similar cell types as observed in existing zebrafish atlases for similar developmental stages as we analyzed in stickleback (Farnsworth, Saunders, and Miller 2020; Wagner et al. 2018). Additionally, we compared the expression of *sox9a* and *sox9b* in our atlas to published descriptions of stickleback *in situ* expression at the same stage (Cresko et al. 2003). Supporting our cluster annotations, we observed expression of both SOX genes in roughly the same cell types as defined by *in situ* hybridization (Figure S2A and S2B; Cresko et al. 2003). Despite our success using a single SMRT cell for ScISOr-Seq for isoform annotation, we would need to sequence additional SMRT cells to have enough data to correlate cell types and specific isoforms.

Since the pooled and merged dataset has higher numbers of cells, genes per cell, and total genes detected than the Ensembl scRNAseq dataset (Figure 2B), assessing the degree to which pooled and merged gene models improved the scRNAseq analysis is complex. For instance, a higher overall expression of a gene in a cluster might be the result of different clustering patterns, more cells retained, and/or more reads counted. We compared the raw counts of specific genes, number of cells expressing these genes, and number of clusters where at least 10% of cells expressed them (Figure 4B). Since Cell Ranger provided a subset of these values on a global scale (Figure 2B), we chose to examine changes in gene counts, cell number, and number of clusters with a case study: the *frizzled* genes, a family of Wnt-pathway signaling molecules expressed in a wide range of tissues (Huang and Klein 2004; Wang et al. 2016). For all *frizzled* genes, we observed increases in raw counts and number of cells expressing them (Figure 4B). Some genes, however, exhibited dramatic increases (*fzd1, fzd4, fzd2,* and *fzd8b*) while others exhibited minor increases (*fzd9b* and *fzd10*). Comparing the gene models of *fzd1* and *fzd9b,* we determined which factors influenced expression detection. Importantly, *fzd1’s* 5’ and 3′ UTR expanded in the merged and pooled annotation, however only the 5’ end of *fzd9b* was extended (Figure 4C and 4D). Based on the read alignments (Figure 4C and 4D), *fzd1’s* increase in counted reads is solely due to the 3′ UTR changes, while *fzd9b*’a slight increase is due to the 5’ end extension. Cluster numbers also varied within the group. While *fzd1, fzd4, fzd2,* and *fzd8b’s* were expressed in more clusters, *fzd9b* and *fzd10* were in one less cell class in the merged and pooled dataset than the Ensembl dataset. We hypothesize that two clusters (containing *fzd9b* and *fzd10*) were combined in the merged and pooled dataset relative to Ensembl. Overall, this gene family comparison illustrates the problem of using an annotation with missing 3′UTRs. If the Ensembl dataset’s *fzd* counts were accepted as their true qualitative and quantitative expression, flawed conclusions would have been made regarding the contributions of each gene in the family.

## SUMMARY

Incomplete 3′UTR annotations can hinder single cell transcriptional profiling studies. In addition to reducing the overall number of genes included in an analysis, systematic differences in preliminary UTR annotations could lead to significant inferential errors. We illustrated in stickleback fish that a minimal ScISOr-Seq dataset, generated concurrently with scRNAseq data, was capable of dramatic improvements in retained read counts (+26.1%). Additionally, we showed that pooling reads from adult bulk Iso-Seq and merging with the existing Ensembl annotations improved reads retained negligibly beyond the models solely resulting from embryonic ScISOr-Seq (+0.8%) suggesting developmental stage of specimens is an important factor when improving annotations. Using the improved annotation for scRNAseq permitted identification of cell types and increased the observed expression of numerous genes. Overall, our work illustrates that ScISOr-Seq is a rapid and cost-effective method to annotate genomes of various organisms for scRNAseq and can improve efficacy of biological inferences.

## METHODS

### Tissue Dissociation to Generate a Pool of Single Cells

We crossed a laboratory line of stickleback originally isolated from Cushman Slough (Oregon) and raised embryos to 70 hours post fertilization (hpf) at 20°C using standard procedures from the Cresko Laboratory Stickleback Facility (Cresko et al. 2004). We euthanized 36 embryos in MS-222 following IACUC approved procedures then dechorionated and deyolked them at room temperature. We limited our embryo dissection to 20 minutes based on Farrell et al. (2018). Following protocols from Bresciani, Broadbridge, and Liu (2018), we dissociated the cells for 6 minutes in 0.25% trypsin in PBS at 30°C, pipetting up and down every 30 seconds, then, stopped the dissociation using 10% FBS DMEM and spun down cells at 400×G for 3min at 4°C. We resuspended cells in 1mL of 0.1% BSA PBS, centrifuged at 400xG for 3min at 4°C, and resuspended in 100ul 0.04% BSA in PBS. At room temperature, we filtered cells through a 40uM cell strainer (Thomas scientific 1181X52) then washed the original tube twice with 100ul of 0.04% BSA in PBS and poured over the same cell strainer.

### Library Preparation

The scRNAseq and ScISOr-Seq samples were prepared and sequenced by the University of Oregon Genomics and Cell Characterization core facility (https://gc3f.uoregon.edu). Samples were diluted to 800 cells/ul using 0.04% BSA in PBS to target 10,000 cells for the 10X Genomics Single Cell 3′ Genome Expression (GEX) mRNA-Seq prep with v3.1 NextGem chemistry. The scRNAseq samples were then sequenced on 1/7^th^ of a single S4 lane on a NovaSeq 6000 platform (Illumina). For the ScISOr-Seq library, 400ng of the amplified single cell cDNA as input for the SMRTbell Express Template Prep Kit 2.0 (P/N 100-938-900) without reamplification. Sample specific barcode (ATATAGCGCGCGTGTG) were added using the Barcoded Adapter Kit 8B—OVERHANG (P/N 101-628-500). The ScISOr-Seq library was sequenced on a single SMRT Cell 8M on the PacBio Sequel II platform using the v4 primer, v2.1 polymerase, 1 hour binding, 30-hour movie, and 2-hour pre-extension time at a loading concentration of 100pM.

### PacBio Data Processing

The University of Oregon Genomics and Cell Characterization core facility (https://gc3f.uoregon.edu) generated “circular consensus” reads for our ScISOr-Seq dataset (ccs -j 39 --min-passes 3 --min-snr 2.5 --min-length 10 --max-length 50000 --min-rq 0.99; v6.6.0) and used lima (-j 39 –isoseq; v2.2.0) to remove a sample specific barcode (ATATAGCGCGCGTGTG) as a part of the PacBio SMRT Analysis software (Supplemental file 6). After barcode clipping, we used a custom ScISOr-Seq processing script (scISOr_Seq_processing.py) that removed the sample primers (3p: CTACACGACGCTCTTCCGATCT; 5p: CCCATGTACTCTGCGTTGATACCACTGCTT), removed and saved cell and UMI barcodes, removed poly(A) tails, then filtered out duplicated reads. This script outputs reads that had the expected presence and orientation of primers and polyA tails for downstream analysis. The script additionally outputs reads without the expected primers in a separate file; however, these other reads were not used for further analysis.

We aligned reads to the stickleback genome (BROAD S1, 104.1 database version) using minimap2 (v2.7; parameters: -ax splice, -uf, -06,24, -B4; Li 2018). We clustered for unique transcripts with collapse_isoforms_by_sam.py script (c=.99, i=.95; Tseng 2021) from cDNA Cupcake (v27.0.0), classified transcripts with the sqanti3_qc.py script from SQANTI3 (v4.2), and refined transcripts with sqanti3_RulesFilter.py script (Tardaguila et al. 2018). We used the sqanti_classification.filtered_lite.gtf that resulted from the refining step (Supplemental file 2) to run Cell Ranger from 10x Genomics (v3.2.0) as described below.

We downloaded the raw data from Naftaly, Pau, and White (2021) from NCBI. This adult Iso-Seq dataset contains gonad, pronephros, brain, and liver reads from both sexes, a total of 16 SMRT cells. We processed reads from this dataset with the following steps: we generated circular consensus reads with ccs (--min-rq=.9; v6.0.0), clipped barcodes listed in Naftaly, Pau, and White (2021) with lima (--isoseq –dump-clips; v2.2.0), removed polyA tails with isoseq3 refine (--require-polyA; v3.4.0-0), and clustered reads with isoseq3 cluster (v3.4.0-0) from the PacBio SMRT Analysis software (Supplemental file 6). For the analysis with solely this data, we aligned reads to the stickleback genome (BROAD S1, 104.1 database version), collapsed transcripts with cDNA Cupcake (v28.0.0), and filtered and classified with SQANTI3 (v4.2) as done with the ScISOr-Seq data. Again, we used the sqanti_classification.filtered_lite.gtf from SQANTI3 (Supplemental file 3) to run Cell Ranger as described below. Due to reduced novel and increased annotated genes relative to Naftaly, Pau, and White (2021), we additionally tried using minimap2-2.15 with -ax splice -uf -C5 -secondary=no to match their parameters; however, we observed negligible changes. We propose that differences in our analyses arise from different earlier processing steps, different versions of stickleback gene models, and an updated version of SQANTI.

We pooled both sets of Iso-Seq reads and merged it with existing Ensembl annotations to create an improved version of the Ensembl annotations. Previously, we created a modified version of the Ensembl annotations where we extended the 3′ UTRs (Supplemental files 7,8) for several marker genes (*tbx16, sox10, sox32*, and *eya1*) and fgf/fgfr genes (*fgfr1a, fgfr1b, fgfrl1a, FGFRL1, fgfr2, fgfr3, fgfr4, fgf3, fgf4, fgf16, fgf17, fgf6, fgf6a, fgf8a,* and *fgf8b)*. Because we lacked ScISOr-Seq reads for *fgf4,* we used this modified gtf for the merging (Supplemental file 8). We completed the processing of the ScISOr-Seq and bulk Iso-Seq data separately as described above then combined the files prior to alignment. We aligned the combined files with minimap2, collapsed transcripts with cDNA cupcake, and refined and classified isoforms with SQANTI3 as explained above.

We merged the sqanti_classification.filtered_lite.gtf resulting from SQANTI3 with the Ensembl annotation using TAMA (Kuo et al. 2020). To prepare the gtf files for TAMA Merge, we converted them to bed files using bedparse gtf2bed (--extraFields gene_id). Then, modified the output with awk to rearrange the columns of the bed file such that the gene id was separated from the transcript id by a semicolon (awk -v OFS=’\t’ ‘{print $1,$2,$3, $13 “;” $4, $5, $6,$7,$8,$9,$10,$11,$12}’). We merged with TAMA’s script tama_merge.py (-s ensembl -cds ensembl -d merge_dup). We applied the ensembl gene names to genes with corresponding IDs using a custom script (tama_associating_ensembl_ids_with_genes.py). For our final gtf file, we converted the output bed file back to a gtf with TAMA’s tama_convert_bed_gtf_ensembl_no_cds.py (Supplemental file 4). We also generated a gtf file specifically for scRNAseq analysis where mitochondrial genes started with “MT” by transforming the bed file with a custom script (tama_associating_ensembl_ids_with_genes_for_scRNAseq.py) and then converting the output back to a gtf in the same fashion as above (Supplemental file 5).

### scRNAseq analysis

We quantified gene counts using 10x Genomics Cell Ranger v3.0.2 (Supplemental file 1). We generated a reference using mkgtf and mkref, retaining protein coding and non-protein coding genes, and then aligned and counted reads using the stickleback genome (BROAD S1, 104.1). We analyzed the counts using the Seurat package (v3.2.3; Stuart et al. 2019) on R (v4.0.2). We retained all cells for the analysis. We normalized the counts using SCTransform and regressed out the mitochondrial genes using the glmGamPoi method (Hafemeister and Satija 2019). Based on the inflection point in the elbow plot of the PCA results, we chose 38 dimensions for generating the UMAP and identifying clusters. We identified cluster identities using three marker gene approaches.

First, we searched for marker genes identified in Farnsworth *et al*. (2020) for specific cell types found in a similar developmental stage from zebrafish (24hpf in zebrafish). Next, we identified markers with Seurat’s FindAllMarkers searching for genes that were positively upregulated in each cluster compared to the other identities, in a minimum of 25% of cells, and a log fold change threshold of 0.25. Finally, we identified markers with Seurat’s FindMarkers searching for genes that were positively upregulated in each cluster versus all the other cells. For the second and third approaches, we searched for the homologous gene in zebrafish and used expression data from zfin to predict which cell types expressed the gene. To quantify *frizzled* gene counts, we used FetchData to isolate raw counts for each gene (*fzd1, fzd4, fzd2, fzd8b*, *fzd9b,* and *fzd10*) and the number of cells expressing the gene. To determine the number of cell clusters each *frizzled* gene was expressed in, we saved a dotplot, subsetted for clusters where the percent of cells expressing the gene was greater than 10%, and then counted the number of clusters left for each gene.

## Acknowledgments

We are grateful for M. Weitzman, D. Turnbull, and J. Sydes from the UO GC3F for assistance with library preparation, sequencing, and initial ScISOr-Seq data processing (ccs and lima). We especially thank C. Small for insightful comments on the analysis, M. Currey for maintaining the stickleback colony, and E.A. Beck for perceptive remarks on the manuscript. We also thank J. Searcy for assisting us at downloading Naftaly, Pau, and White (2021)’s raw data from google cloud server. This work was funded by a PacBio local SMRT grant (ScISOr-Seq library preparation and sequencing), This work was funded by National Science Foundation grant OPP-2015301 and Oregon Research Excellence Funds (to WAC). HMH was supported by the Genetics Training Program (NIH T32GM007413) at UO.

## Data Accessibility

All raw sequencing data associated with this study will be available upon peer-reviewed publication via the NCBI Sequencing Read Archive (SRA), under BioProject PRJXXXXXXXX. Assembly, annotation, and summary files not in the Supplementary Information will be available upon peer-reviewed publication in the Dryad repository under Accession XXXXXXXX. All scripts can be found in the github repository (https://github.com/hopehealey/scISOseq_processing).

## Supplemental Figures

**Figure supplement 1:**
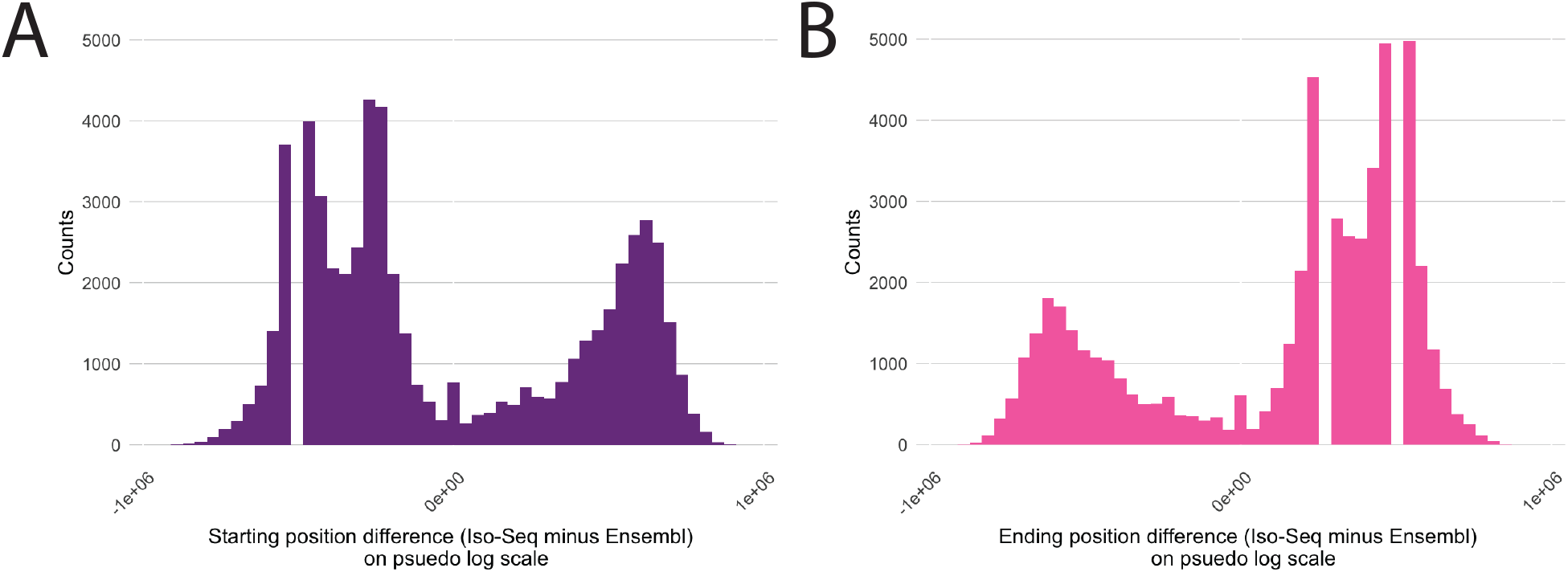
By comparing the median difference in starting or ending position between the ensembl annotations and pooled Iso-seq annotations, it shows that bulk Iso-Seq and ScISOr-Seq reads overall lead to earlier starting positions (median=−62, A) and extended ending positions (median=62, B). We plotted this on a pseudo log scale to account for the negative values.

**Figure supplement 2:**
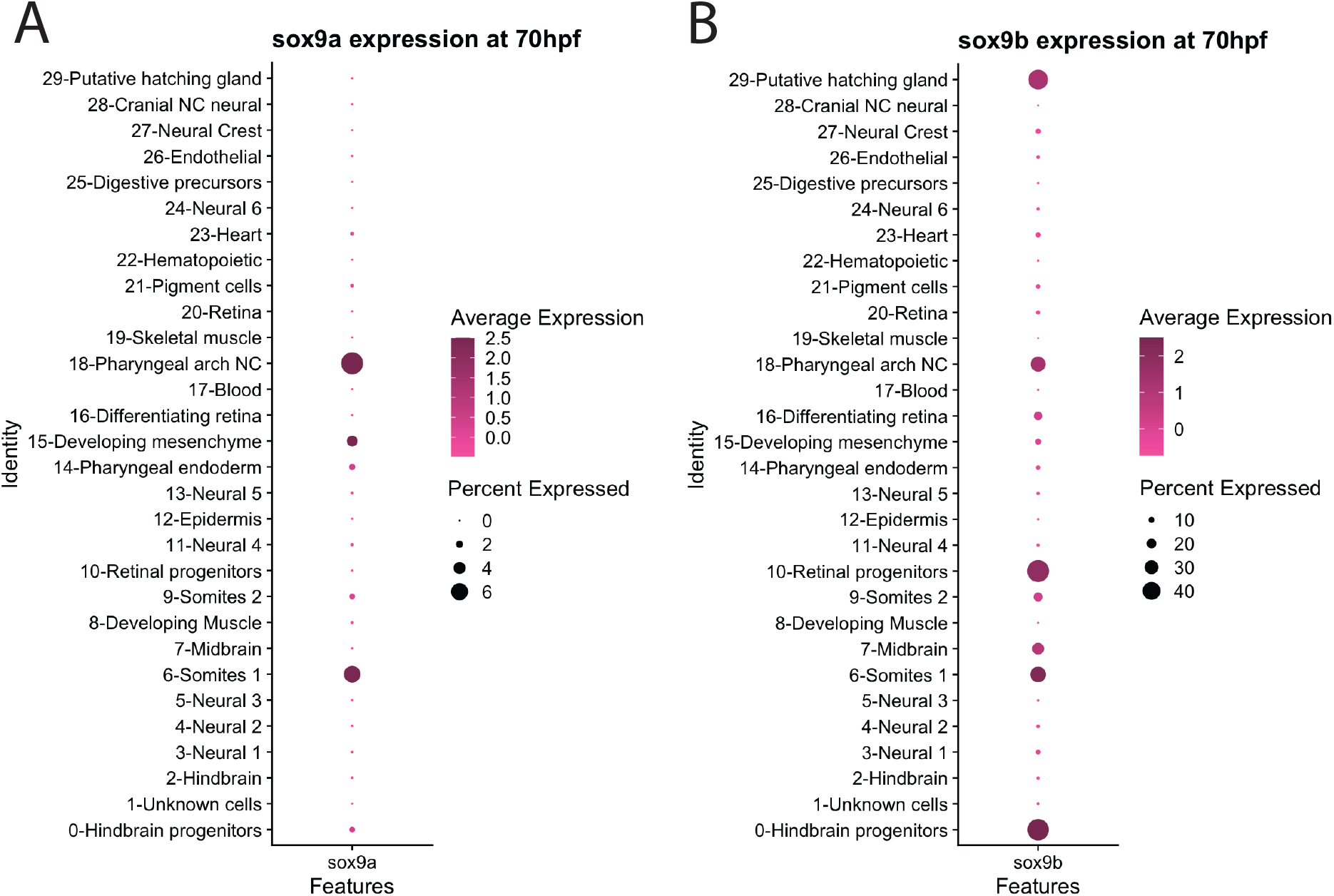
Expression patterns of *sox9a* and *sox9b* recapitulate expression seen in Cresko *et al.* 2003. The two paralogs are plotted on separate Dot plots due to an order of magnitude difference in expression. A) as observed in Cresko *et al.* 2003, *sox9a* is expressed in the pharyngeal arches, mesenchyme, and endoderm as well as the somites; however, we did not observe any *sox9a* expression in the forebrain. B) We found *sox9b* expression in the pharyngeal arches, retina, somites, hindbrain, and midbrain similar to Cresko *et al.* 2003, though our dataset lacked forebrain *sox9b*.

## Supplementary file captions

**Supplementalfile1_CellRangerOutputInformation.xlsx:** A table containing detailed Cell Ranger results for each annotation that was used.

**Supplementalfile2_Annotation1_scISOrSeq.gtf**: The genome annotations that originated from the scISOrSeq sequencing reads and was produced from SQANTI3.

**Supplementalfile3_Annotation2_Bulk_Iso_Seq.gtf**: The genome annotations that originated from the bulk Iso-Seq sequencing reads from Naftaly et al. (2021) and was produced from SQANTI3.

**Supplementalfile4_Annotation3_Pooled_Iso_Seqs_merged_with_ensembl.gtf**: The genome annotations from ensembl (BROAD S1, 104.1) merged with the gtf file produced by SQANTI3′s file that originated from the bulk Iso-Seq and scISOrSeq data.

**Supplementalfile5_Annotation4_Pooled_Iso_Seqs_merged_with_ensembl_for_cellranger.gtf**: The genome annotations from ensembl (BROAD S1, 104.1) merged with the gtf file produced by SQANTI3′s file that originated from the bulk Iso-Seq and scISOrSeq data. The file is the same as the annotation 4 file with the exception of mitochondrial genes which have gene names that start with MT.

**Supplementalfile6_IsoSeq_Processing_Data.xlsx**: The processing information (reads retained and lost at each step) for the Bulk Iso-seq and ScISOr-Seq datasets.

**Supplementalfile7_EnsemblModified_manual_changes.xlsx**: The 3′ UTR modifications made to the ensembl genome for select marker genes and *fgf*/*fgfr* genes.

**Supplementalfile8_Annotation5_ensembl_modified.gtf**: The genome annotations from ensembl (BROAD S1, 104.1) with 3′ UTRs extended in select marker genes and *fgf*/*fgfr* genes as described in **EnsemblModified_manual_changes.xlsx**.

## Notes

### Competing Interest Statement

The authors have declared no competing interest.

